# Methods for estimating location from spatial patterns in species composition: a fishing location case study

**DOI:** 10.64898/2025.11.30.691468

**Authors:** James A. Smith

**Affiliations:** Port Stephens Fisheries Institute, New South Wales Department of Primary Industries and Regional Development, Locked Bag 1, Nelson Bay, NSW 2315, Australia

**Keywords:** species distribution modelling, joint species model, commercial fishing, survey location, location uncertainty

## Abstract

The location at which a species assemblage is observed can be uncertain, for example in fisheries where catch composition is recorded accurately but location data may be coarse or inaccurate. In such cases, spatial signals in observed species compositions can help identify and refine uncertain locations.

This study explored three approaches for location estimation using species composition data: 1) a hierarchical species distribution model (SDM) that jointly estimates species distributions and location uncertainty, 2) an inverse prediction method that uses a fitted SDM to identify the most likely location given new species data, and 3) the direct modeling of location as the response variable. Each approach requires a subset of observations with accurate locations to quantify the spatial patterns in species distributions.

All three methods were useful and reasonably accurate: for the simulated data the average distance error (distance to true location) was 15% the size of the domain, and for the real data this error was 20-100 km. When only some locations are uncertain, the hierarchical approach is valuable due to its integrated estimation of location and species parameters. When many locations are uncertain, the other approaches seem more suitable. Inverse prediction is ideal when prior information is available to constrain the location estimation. Direct modelling (although causally spurious) is well suited to large datasets, with multivariate random forests often providing the most accurate location estimates.

These methods can enhance spatial data quality in ecological monitoring and help develop tools for improving inaccurate or deliberately misreported fishing locations.

## 1. Introduction

In many ecological datasets, the location at which species compositions were observed may be unknown or recorded inaccurately, while the species composition data are diverse and informative. This is especially the case for commercial fishing – the location at which a catch is made is often uncertain. Reasons for this include a lack of reporting requirements, spatially coarse reporting requirements, deliberately or accidentally inaccurate reporting, and illegal fishing (Bastardie et al 2010, Sampson 2011, Mangi et al 2015, Emery et al 2019, Brown et al 2021, Watson et al 2023). Marine fisheries in New South Wales (NSW) Australia, typically report location of catches in 0.1 degree cells (NSW DPIRD 2025), but there is anecdotal evidence that cells are sometimes recorded incorrectly, and catches were historically reported in coarser 1 degree cells. Accurate locations are important for various management and monitoring objectives, so refining these locations is of great value.

Accurate fishing locations can be sourced using vessel tracking devices (such as vessel monitoring systems, VMS) and track decoding algorithms (Lee et al 2010, Russo et al 2018), but not all fisheries or vessels use these devices. Accurate locations can also be sourced from scientific observer surveys but these can be occasional and cover a small portion of the fishing effort (Yin et al 2024). Less explored is using the inherent spatial signal in multispecies catches to identify and refine the location of fishing (Russo et al 2016).

The foundation of this idea is that caught species have unique spatial associations and habitat preferences, and these inherent spatial distributions will leave a signal in the sampled species (Watson et al 2023). This is the same idea as species distribution modelling, which identifies these spatial distributions typically by regressing species abundance (including from catch data) against spatio-temporal and environmental predictors (Guisan and Zimmerman 2010, Smith and Johnson 2024). Estimating catch location is similar, except that now location becomes a variable to estimate rather than one assumed known. Although numerous studies include location as a predictor when estimating the distribution of fishing or catch (Brodie et al 2019), it is a related but different task to estimate location based on patterns in catch.

There are a few approaches for using catch composition to help refine fishing location, and in this study they all rely on a subset of locations being well known. An example is occasional scientific observer surveys of commercial fishing – the observed subset accurately measures location and can be used to refine the unobserved uncertain locations. The three approaches evaluated in this article are (summarised in Table 1): 1) model catch (the outcome) as a function of location, but estimate location error for uncertain locations by leveraging the information from the accurate locations in a single hierarchical model, called the *location error* approach; 2) model catch as a function of location for the accurate locations only, then invert the model to predict location given catch composition for the uncertain subset, called the *inverse prediction* approach; and 3) model location (the outcome) as a function of catch composition for the accurate locations, and then simply predict location for the uncertain subset, called the *location as response* approach.

**Table 1:**
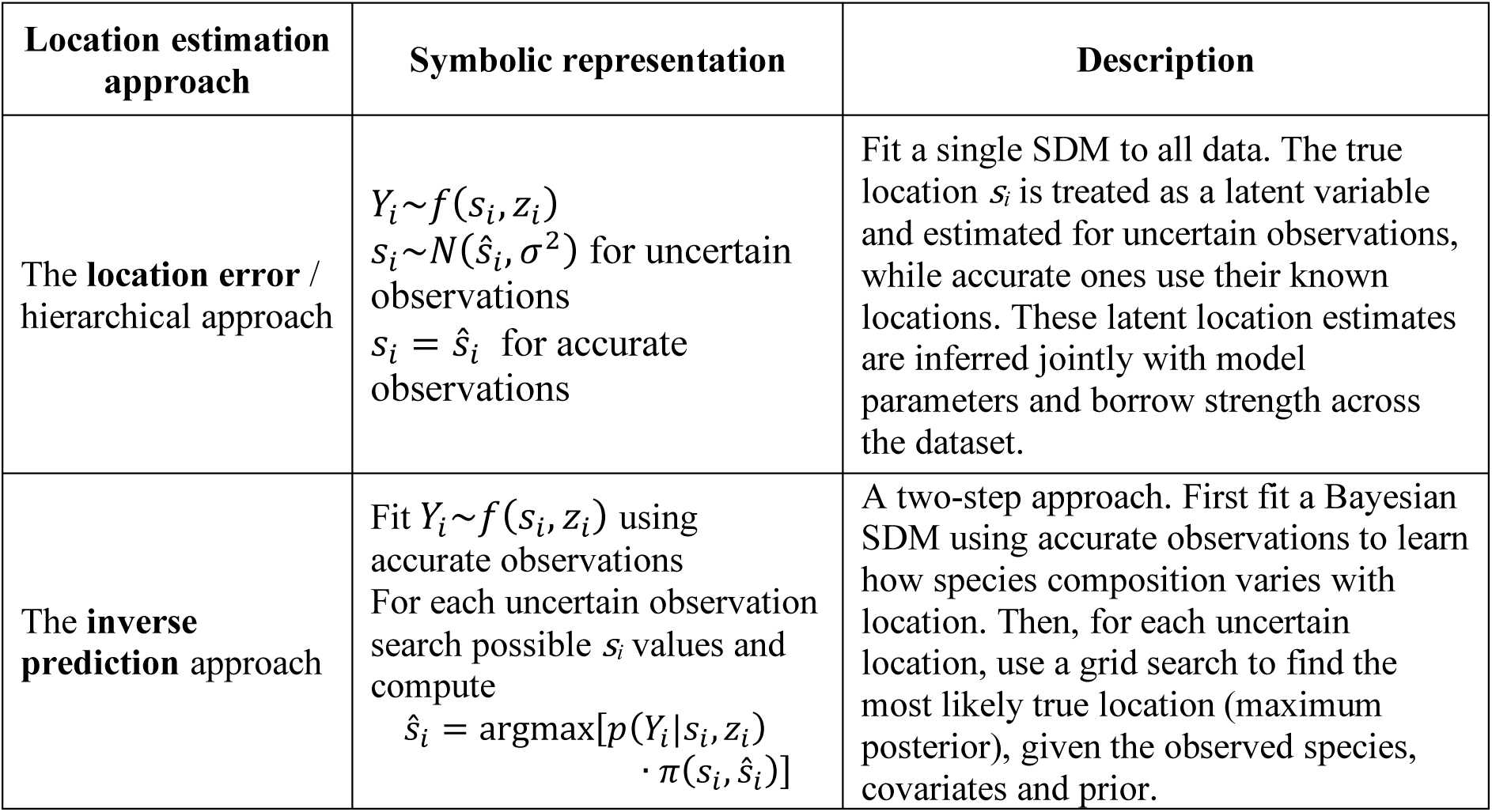
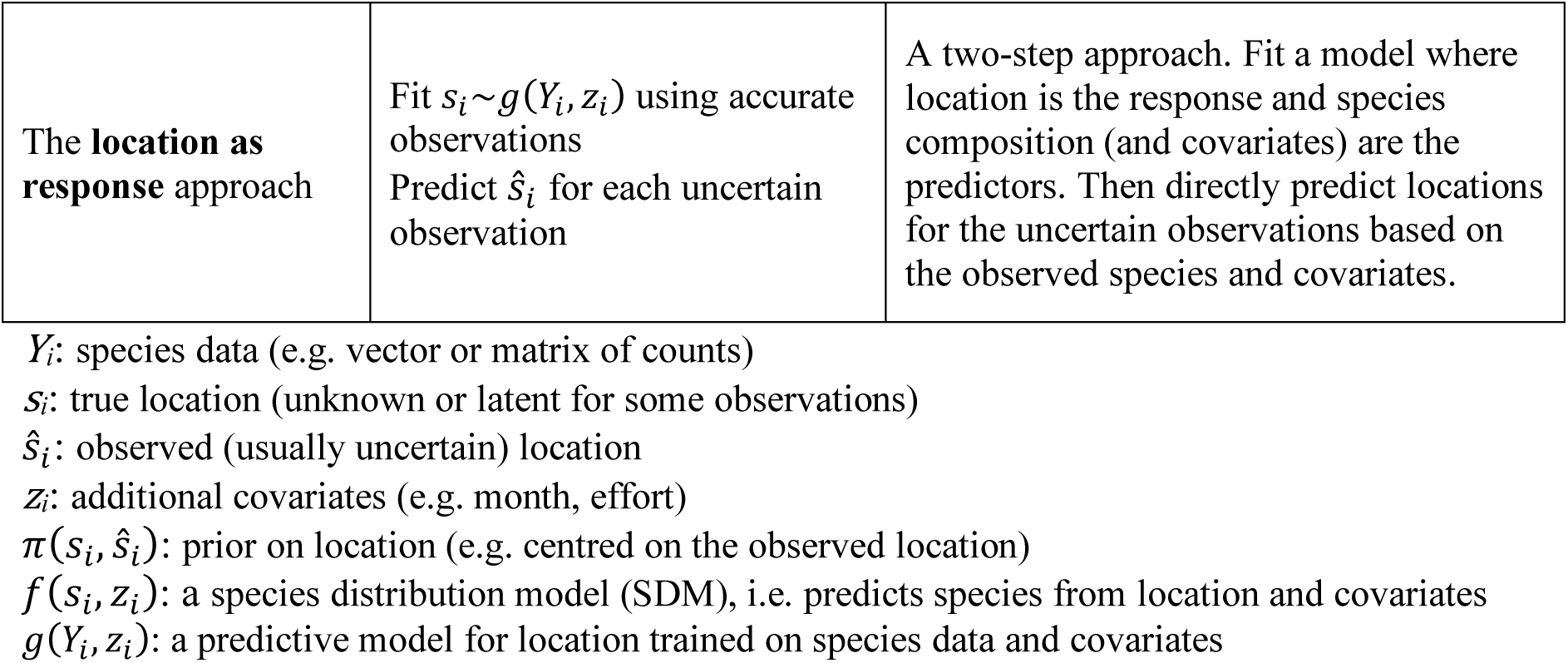
A summary of how each modelling approach estimates uncertain (or unknown) sampling locations. More detailed equations are provided in the methods.

Each approach has apparent advantages and disadvantages, and this article presents and evaluates them using simulated and actual fishing catch data. The goal is to provide a conceptual and numerical workflow for similar analyses that desire to estimate, refine, or check the location of sampling events using multispecies observations. Although this objective is relevant to numerous types of ecological data and applications, the focus here is on estimating fishing locations.

## 2. Methods

Three approaches were tested using both simulated and real data (Table 1). Each approach and its model implementation are described below, followed by details of the data simulation process, performance metrics, and the scenarios used for evaluation. The real catch data used to assess accuracy in a real-world context are then outlined. These approaches can refine uncertain locations and estimate unknown ones, but for brevity locations are described below as “uncertain”.

### 2.1. The location error (hierarchical) approach

This approach uses a single joint hierarchical Bayesian model which treats the true location of each observation as a latent variable. Observed locations are modeled as noisy measurements of the true location, with uncertainty levels that vary by observation, i.e. accurate locations have no uncertainty. The goal is to estimate posterior distributions over these latent locations jointly with other parameters, such as species-environment relationships. This allows all available data (accurate and uncertain) to inform spatial structure while accounting for location uncertainty. The advantage of this approach is that an integrated model is used to model the relationships between species and location for accurate records to update the location of uncertain records. An integrated model has neat statistical properties and jointly estimated uncertainty. The main disadvantage is that the proportion of accurate to uncertain records will influence the accuracy of the updated estimates, and when there are many uncertain records this approach may be inaccurate and computationally slow.

This approach treats the true bivariate locations *s*_*i*_ = (*s*_*i*,*x*_, *s*_*i*,*y*_) as latent variables (i.e. unobserved) for observations with uncertain location, while treating the accurate locations as exactly known. The observed locations *ŝ*_*i*_ = (*ŝ*_*i*,*x*_, *ŝ*_*i*,*y*_) are considered noisy measurements of *si*. A species distribution model (SDM) models species data as:

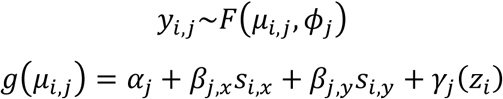

where *F* is the species-specific probability distribution (e.g. negative binomial), *g* is a link function, *y*_*i*,*j*_is abundance of species *j* in observation *i*, *z*_*i*_ are additional covariates (e.g. month), 𝜇_*i*,*j*_ is the expected mean abundance, 𝜙_*j*_are any additional shape or dispersion parameters, 𝛼_*j*_is the intercept term, 𝛽_*j*,*x*_and 𝛽_*j*,*y*_are the spatial effects, and 𝛾_*j*_ are additional covariate effects. The observation error component is defined for observations with uncertain locations as:

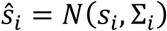

where Σ_*i*_ is the prior covariance matrix, which in this case assumes independent standard deviations for each axis (𝜎_*si*,*x*_and 𝜎_*si*,*y*_) which are specified. For observations with accurate locations we assume *ŝ*_*i*_ = *s*_*i*_. The Bayesian posterior distribution which combines the measurement model and the observation model (the SDM) is thus:

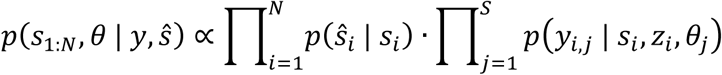

This is a hierarchical model because it includes observation error and data models, and a joint (or multivariate) model because it estimates species relationships simultaneously, although it does not (but could) estimate dependence among species such as their residual correlations (Warton et al 2015). Location uncertainty was quantified using the full posterior distribution of the location estimates. For each uncertain observation, marginal 90% credible intervals were calculated for both spatial coordinates for model comparisons and mapping.

### 2.2. The inverse prediction approach

This approach flips the usual predictive task – rather than predicting species from location, we predict location from species. The idea is to fit a Bayesian SDM using accurate locations, invert the model (i.e. use the SDM as a likelihood function) to predict the most likely locations for a new set of species composition data. This inverted prediction combines the SDM with a prior over possible locations to estimate a posterior distribution. The advantages of this approach are that a single fitted model can be used to predict many uncertain locations, that a variety of priors can be placed on location variables (Fig. 1), and that additional covariates can be accounted for in model fitting and prediction. Having a prior for this problem is very useful, e.g. catch records may be inaccurate but still informative within a broader area and an informative prior can refine the search area. Alternatively, an uninformative prior can be used to identify entirely unknown locations. The disadvantages of the inverse approach are that the joint likelihood may be complex and challenging to maximise, and that a grid search (or optimisation) must be done for each data record which can be slow.

**Fig. 1.**
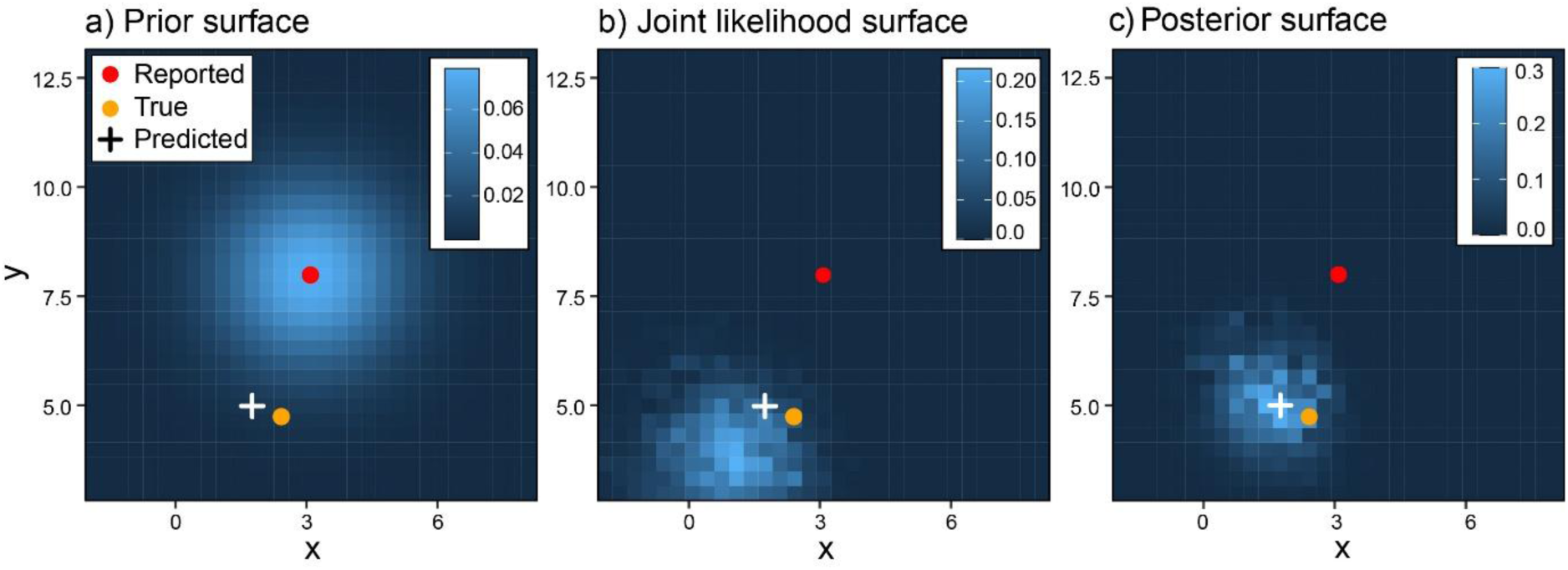
The likelihood surfaces for the inverse prediction approach. For an example observation this shows the true location of catch (orange dot) and uncertain reported location (red dot). The prior (a) is centred on the uncertain; the joint likelihood (b) is how likely each location is given the observed species composition and fitted SDM; and the posterior is the combination of the prior and joint likelihood (c) which is used to predict the most likely location of fishing (cross).

This approach assumes that for data with uncertain spatial information, the true (but unknown) bivariate location is *s* = (*s*_*x*_, *s*_*y*_), and the observed species composition is *y* = (*y*_1_, *y*_2_ …, *y*_𝑁_), i.e. counts of *N* species. A prior distribution is placed over the true location *s*, denoted 𝜋(*s*), and an SDM is fitted on accurate data, potentially including other covariates *z* (e.g. month), to compute the likelihood of observing *y* at location *s*. The SDM provides:

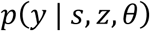

where *θ* are the parameters of the fitted model. For example, if the SDM is a multivariate generalized linear model with independent species likelihoods, the joint likelihood is:

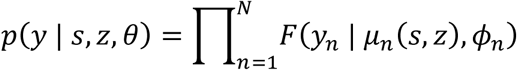

where *F* is the species-specific probability distribution (e.g. negative binomial), 𝜇_𝑛_(*s*, *z*) is the predicted mean abundance of species *n* at location *s*, and 𝜙_𝑛_are any additional shape or dispersion parameters. This likelihood can be generalised to numerous similar forms depending on the structure of the species distribution model, e.g. a joint species model estimating residual species correlations (Warton et al 2015) would not have independent species likelihoods.

The Bayesian posterior over locations is:

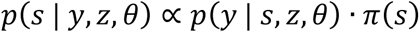

The prior 𝜋(*s*) can take numerous forms, but a reasonable choice in a bivariate normal centered at the approximate reported location *s*_0_(Fig. 1):

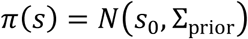

where Σ_prior_ is the prior covariance matrix, here assuming independent standard deviations for each axis (𝜎_*sx*_and 𝜎_*sy*_). This formulation allows *sx* and *sy* to represent geographic coordinates (e.g. longitude and latitude) or other combinations such as depth and latitude.

Rather than optimizing 𝑝(*s* | *y*, *z*, 𝜃) directly, which may be error-prone for complex likelihood surfaces, it is evaluated over a discrete grid of candidate locations (*s*_1_, *s*_2_ …, *s*_𝐾_). For each grid point, we calculate:

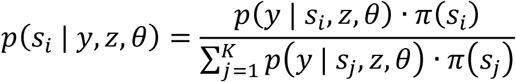

The grid point *s*_*i*_ with the highest posterior probability is taken as the maximum a posteriori estimate (MAP), i.e. the most likely location of catch given the observed data (Fig. 1).

To quantify uncertainty, 90% marginal credible intervals are approximated for each axis by first computing the marginal distributions (summing posterior probabilities over all y-values for each x-coordinate, and vice versa) and then identifying the 5th and 95th percentiles of the resulting cumulative distribution functions.

### 2.3. The location as response approach

The idea of this approach is that we fit models with species composition as predictors and location as the response (e.g. Watson et al 2023). The accurate locations are used to describe how location can be identified by values of each species and additional covariates, then we simply predict location for the uncertain locations based on their observed species and additional covariates. The advantage of this approach is its simplicity and straightforward prediction. A key disadvantage is that the causal flow is illogical, because the model assumes that location responds to species when in reality species respond to location. While this logic may not matter for a prediction task it could still lead to some spurious predictions, e.g. the location–abundance relationship for a species that has similar abundance everywhere would be highly unstable. Another disadvantage is that modelling species jointly (e.g. estimating their residual correlation) is not feasible which could reduce the predictive power.

This approach does not benefit from a Bayesian framework, so two flexible model types are tested: a multivariate generalized additive model (GAM) and a multivariate random forest. The multivariate GAM estimates the residual correlation among location variables using a multivariate normal distribution (Wood 2017). This allows for joint prediction and correlation of the location variables, which means the uncertainty of a predicted location can be elliptical. Random forests do not assume a particular parametric form (Breiman 2001), and will learn flexible relationships between species composition and location (Smith and Johnson 2024). The multivariate random forest incorporates all response variables in the splitting rule (the process used to grow trees) and can adjust this rule based on correlation among responses (Ishwaran et al 2021) which can improve model fit (Segal and Xiao 2011). Uncertainty from the random forest predictions were derived from the variation among individual trees in the forest, which are themselves trained on bootstrapped samples of the data (Breiman 2001).

For the multivariate GAM, we model each dimension of the true location *s* as a smooth function *f* of the observed *N* species *y* and additional covariates *z*:

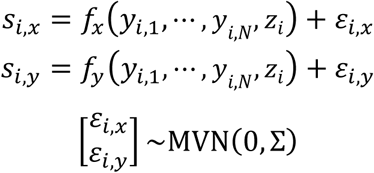

where Σ is the 2 × 2 residual covariance matrix. No equation is shown for the random forest because it does not have a classically interpretable model.

### 2.4. Simulated and real data

To compare the use and performance of these three approaches, multi-species abundance data were simulated using a custom function (Fig. 2). This function generates counts of *n* species (here, *n* = 5) across bivariate space representing, say, longitude and latitude. Each species has a specified mean abundance and dispersion (used in a negative binomial distribution), and specified preferences for the two spatial dimensions and month, customised by a preference magnitude and noise. For simplicity linear preferences were specified on the scale of the linear predictor. The preference for month was included to demonstrate the addition of other covariates. The function operates by first generating a random set of locations, and using the above preferences to calculate the expected abundance (plus noise) of each species at each location. The locations are used exactly for the ‘accurate’ observations, and a specified level of noise is added to the locations for the ‘uncertain’ observations. Stochasticity exists in the addition of noise and the sampling of the negative binomial distribution, so for this reason repetition was included in the scenario testing (see below).

**Fig. 2.**
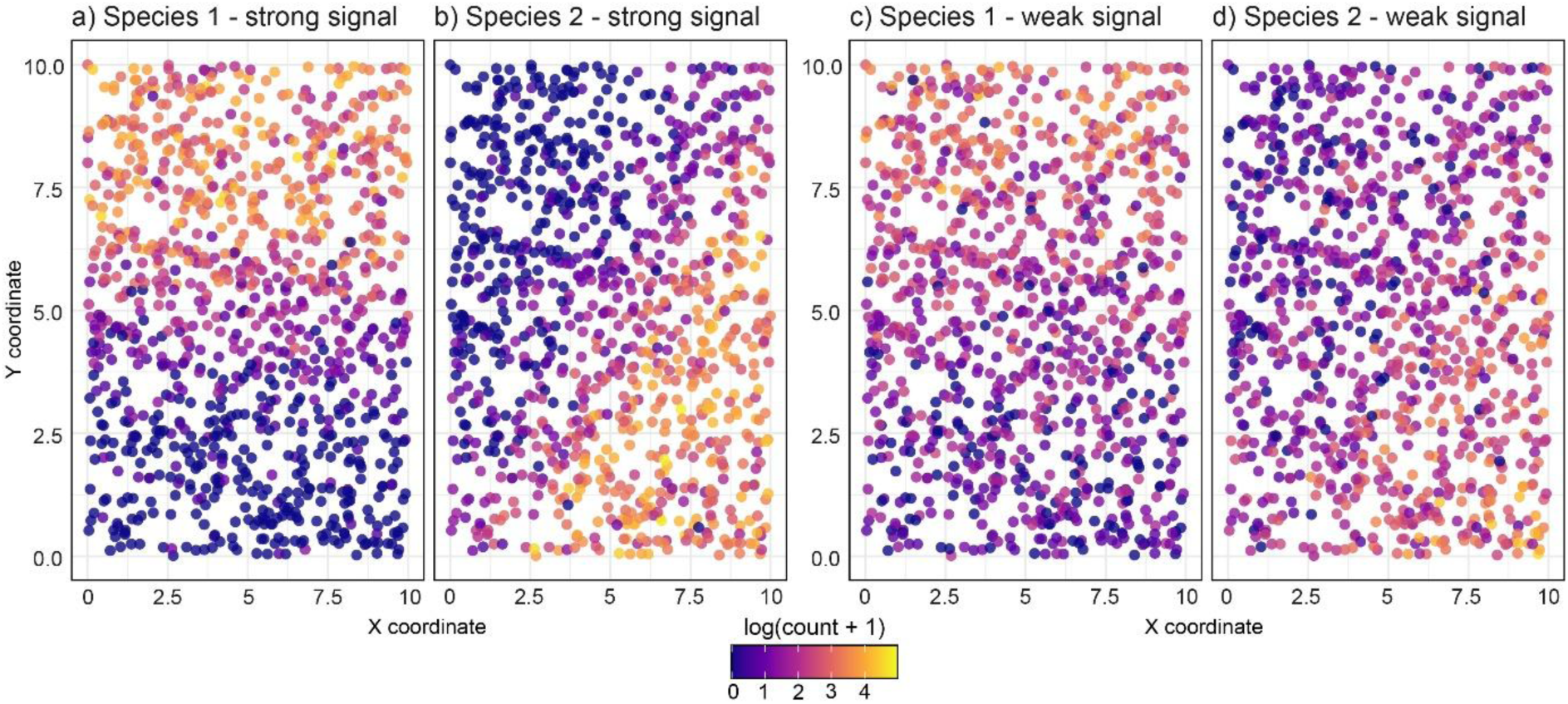
Simulated catches. Catches are counts (n = 1000) for two simulated species with strong spatial signals (a-b) or weaker spatial signals (c-d). An uncertain location will be estimated with higher accuracy when the signal is stronger.

Real multivariate count data was sourced from an ocean prawn trawl in New South Wales, Australia. These data come from a scientific observer survey, so the locations of catch are exactly known (Johnson and Barnes 2023). Nine species were selected that were both relatively common and showed reasonable predictive power in an SDM (Smith and Johnson 2024). These data come from the east coast of Australia, which has a north–south orientation, and from a narrow band of the continental shelf. For this reason, the locations are informative when defined using longitude and latitude and when using bottom depth and latitude.

### 2.5. Tested scenarios

Six scenarios and all three modelling approaches were tested using the simulated data, and two scenarios and two approaches for the real data. These scenarios are summarised in Table 2. In general, the goal for the simulated data was to refine uncertain locations, and the goal for the real data was to estimate unknown locations (unknown within the typical fishing grounds). The scenarios for the simulated data explored how model performance was affected by: 1) the strength of the spatial preferences (Fig. 2); 2) the amount of noise added to locations; 3) the strength of the Bayesian priors (too strict and it may unfairly bias the estimates); and 4) the absolute and relative number of accurate vs uncertain observations. The scenarios for the real data used only the inverse prediction and random forest location as response approaches (based on results from the simulated data), and compared location predictions when defined as longitude–latitude or as depth–latitude.

**Table 2.** Summary of the scenarios tested for the simulated and real multi-species count data. ‘Accurate:Uncertain’ represents the number of observations of each type, which is especially relevant to the location error approach. ^1^ Priors are not relevant to the location as response approach. ^2^ This represents an 85:15 train:test split.

| **Simulated data (5 species)** |  |  |  |  |
| --- | --- | --- | --- | --- |
| **Scenario** | **Accurate:**<br>**Uncertain** | **Species**<br>**signal** | **Location**<br>**error** | **Location**<br>**priors<sup>1</sup>** |
| **Scenario 1:** Base | 300:300 | Strong | Moderate | Informative |
| **Scenario 2:** Weaker spatial signal | 300:300 | Weak | Moderate | Informative |
| **Scenario 3:** Weaker signal and inaccurate location estimates | 300:300 | Weak | High | Informative |
| **Scenario 4:** As above but less strict priors | 300:300 | Weak | High | Weaker |
| **Scenario 5:** Fewer accurate data | 100:1000 | Strong | Moderate | Informative |
| **Scenario 6:** Minority accurate data | 300:3000 | Strong | Moderate | Informative |
| **Real data (9 species)** |  |  |  |  |
| **Scenario** | **Accurate:**<br>**Uncertain<sup>2</sup>** | **Location**<br>**coordinates** |  | **Location**<br>**priors<sup>1</sup>** |
| **Scenario 7:** Geographic coordinates | 1130:200 | Longitude and Latitude (degrees) |  | Weak |
| **Scenario 8:** Depth and latitude | 1130:200 | Depth (m) and Latitude (degrees) |  | Weak |

### 2.6. Model implementation

The location error (hierarchical) approach was modelled using Stan (Carpenter et al 2017) implemented in R (v4.4.1, R Core Team 2024) using rstan (Stan Development Team 2024). A joint modelling framework was used so that all species shared the same estimated locations, but their residual correlations were not estimated. For the simulated data, 4 chains of 4000 iterations were used with 2000 iterations as warm up. The ‘adapt delta’ value was increased to 0.9, which reduces the step size between samples, to remove some divergent transitions. A negative binomial distribution and log link function were specified, as were linear terms for the location variables, to match how the data were generated. The priors were: species intercepts: Normal(1, 1); spatial effects: Normal(0, 0.5); month effect: Normal(0, 0.5); dispersion: Exponential(1). Model suitability was confirmed using the ratio of effective sample size to total sample size (ESS ratio), the scale reduction statistic, and a visual inspection of trace and density plots.

The inverse prediction approach was modelled using the ‘brms’ package (Bürkner 2017) which leverages Stan in R, and was used due to its more user-friendly interface and easier manipulation. As in the hierarchical approach, the models were fit using a joint framework and used the same variable structure and validation process. The location priors are used in the inverse prediction stage, and can be either informative for location refinement (simulated data scenario, Table 2), or uninformative for location identification (real data scenario, Table 2). In the real data scenarios, smoothers were used for the location coordinates to increase model flexibility. These were either isotropic thin plate smoothers for longitude and latitude, i.e. `s(lon, lat, k=20)`, or anisotropic tensor product smoothers for depth and latitude, i.e. `t2(depth, lat, k=20)`. Also included were month and log-transformed area trawled as additional covariates, the latter required to account for differences in sampling effort among observations. The real data required increasing adapt delta to 0.95 to reduce divergent transitions.

The multivariate GAM was fit using the ‘mgcv’ package (Wood 2017). Thin plate smoothers were used for the species covariates. GAM prediction uncertainty was quantified using unconditional standard errors (which include smoothing parameter uncertainty) combined with residual variance to provide full 90% prediction intervals, improving comparability with the Bayesian approaches.

The multivariate random forest was fit using the randomForestSRC package (Ishwaran and Kogalur 2025). For the simulated data scenarios 1000 trees were used, the number of variables randomly sampled at each split (‘mtry’) was 2, and node size was 3; mtry was increased to 3 for the real data scenarios due to the larger number of covariates. Prediction uncertainty was quantified using empirical standard deviations from the bootstrap distribution across trees to provide 90% prediction intervals.

### 2.7. Measuring performance

Performance was measured using two metrics: distance error and percentage of locations improved. These metrics require knowing the true location, which was the case for the simulated and real data scenarios. Distance error was measured as the mean Euclidean distance between true locations and the reported or predicted locations. The percentage of locations improved is simply the percentage of predicted locations that are more accurate than the reported locations. Due to the challenge of deriving comparable uncertainty estimates across approaches, uncertainty is typically evaluated only across scenarios within an approach. For the real data scenario defining location using depth and latitude, we also examined the accuracy of the depth estimates and how these translated when mapped using geographic coordinates.

## 3. Results

### 3.1. Simulated data

All three approaches worked as intended and could estimate location given the simulated catch composition. The estimates and their uncertainty varied considerably among approaches, reflecting their different statistical foundations and their implementation (Fig. 3). When species had a strong spatial signal (Scenario 1), all methods reduced the spatial error (the distance to the true location) by 40-50%, and improved the accuracy of 80-90% of reported locations (Fig. 4a). The location as response approach using a random forest reduced this error the most, but the location error and inverse prediction approaches improved the most locations. However, when the species spatial signal is weaker (Scenario 2), all methods performed worse, with only a 20% reduction in error, and the two location as response models performed the worst (Fig. 4b).

**Fig. 3.**
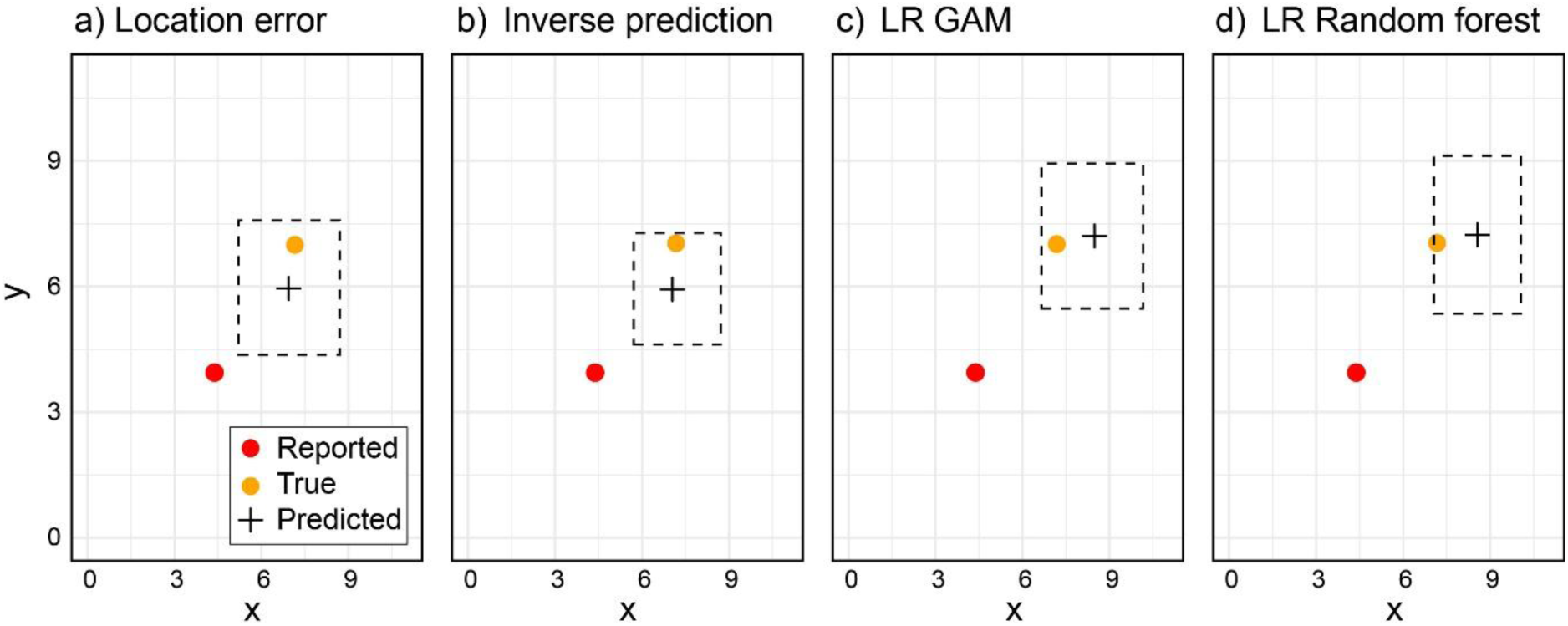
‘Maps’ showing the estimated uncertain location by the four models for an example simulated observation. The spatial dimensions are arbitrary (x and y; the same as in Fig. 2). The red dot shows the reported uncertain location, the orange dot shows the true location, and the cross shows the predicted location (point estimate or MAP). The dotted rectangle shows the approximate 90% marginal confidence bounds, which differ in derivation among models. ‘LR’ indicates the location as response approach.

**Fig. 4.**
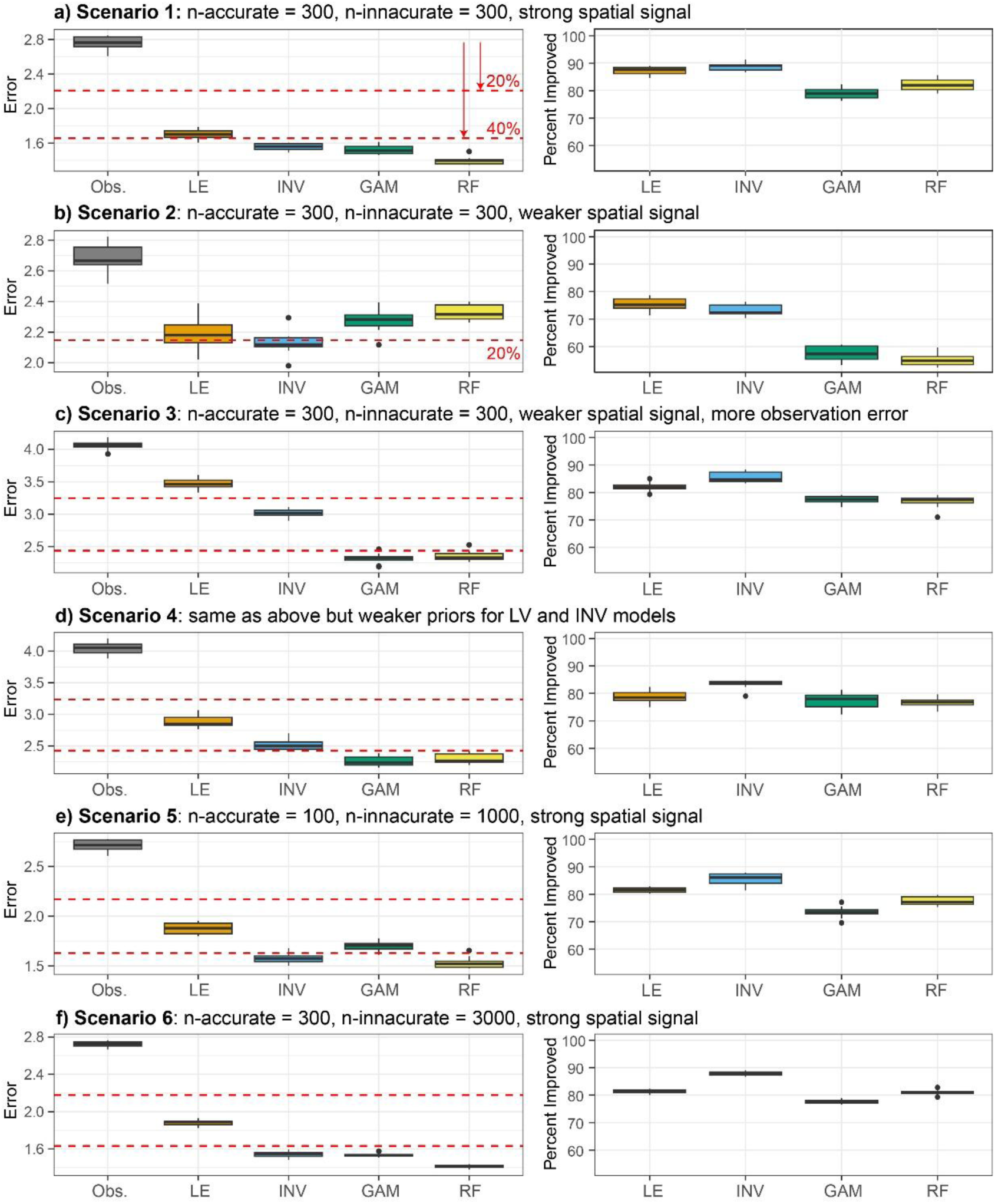
Comparison of model performance for the six simulated scenarios (a-f). The left column shows the mean spatial error (Euclidean root mean square error) between the true simulated locations and the inaccurate observed locations (Obs.) or model predicted locations (LE = location error, INV = inverse prediction, GAM = GAM location as response, and RF = random forest location as response). The mean error for the four models is always lower than the observed error, which shows an improvement in location accuracy under every scenario. The red dashed reference lines show a 20% and 40% decline in spatial error, relative to reported. The right column shows the percentage of predicted locations that were more accurate than the reported locations. Each boxplot contains n = 10 replicates which incorporate stochasticity in the data generation and model fitting processes.

When additional observation error is added (Scenario 3), i.e. the reported locations are farther from the true locations, the two Bayesian approaches performed poorly (Fig. 4c). This was due to disinformation from the prior, and only when the prior was relaxed (Scenario 4) and more likely to include the true location did their performance improve (Fig. 4d). When the number of accurate locations is small relative to the number of uncertain locations to be estimated (Scenarios 5 and 6) we see that the location error approach performed worse (Fig. 4e), and this seems unrelated to the absolute number of accurate observations (Fig. 4f vs 4e).

### 3.2. Real data

A mapped estimated location of catch for an example observation is shown in Fig. 5a (Scenario 7). The true location of catch was in the northern area, and the inverse prediction approach identified a location 85 km away, with large latitudinal marginal uncertainty that encompassed the true location. Across all test observations, the inverse prediction approach had a median distance error of 97 km but with some locations > 500 km from the true location (Fig. 5c). However, the location as response random forest model was much more accurate, with a median distance error of only 21 km (Fig. 5d). If we take the same example observation in Fig. 5a but define location using latitude and depth (Scenario 8) we get the map in Fig. 5b. The latitudinal estimate has shifted north, but there is broad uncertainty in the depth at which the catch occurred. If we map these data on geographic coordinates, we see an estimated location 36 km away and in shallower depths than the true location (Fig. 5e).

**Fig. 5.**
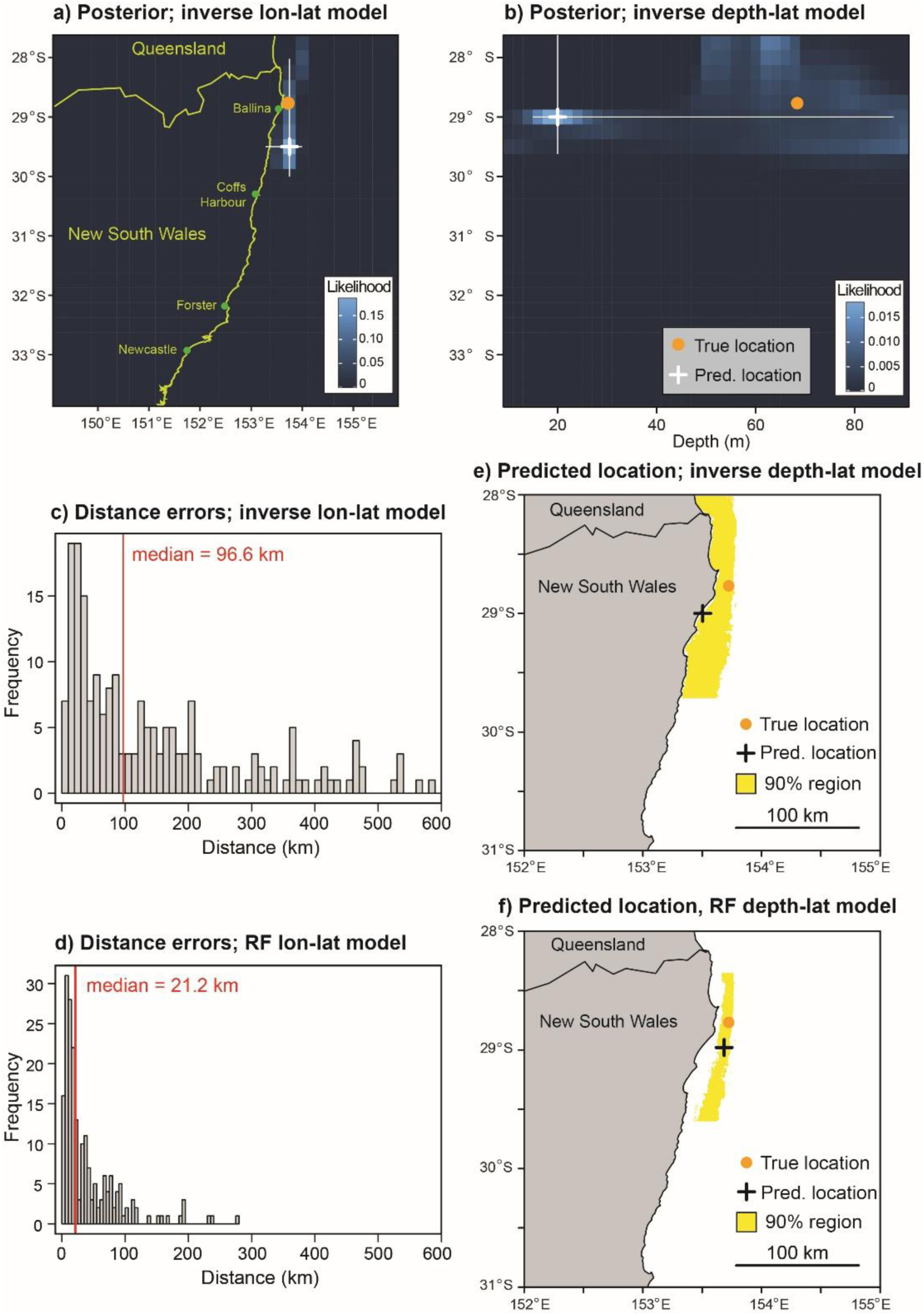
Summary of the scenarios using real prawn trawl data. a-b) For an example observation, the posterior likelihood surfaces for the inverse model that identify location using longitude and latitude (a) or depth and latitude (b). c-d) Histograms of distances (km) between true and predicted locations for a test data set, for the inverse (c) or random forest (RF) model (d) that identify location using longitude and latitude. e-f) Maps of the true and predicted locations for the inverse (e) or random forest model (f) that identify location using depth and latitude; the yellow area is the 90% confidence area for the predicted location. The same example observation is used in panels a, b, e, and f.

Again, the location as response approach was more accurate, with the random forest having a more accurate depth and smaller region of uncertainty (Fig. 5f). Across all test samples, the random forest was on average within 6 m of the true depth, and the inverse prediction model within 17 m (the true fishing locations tend to occur within a 60 m depth range).

## 4. Discussion

The success of using species compositions to estimate the sampling location depends on the strength of the spatial signal in species distributions relative to the spatial error in reported locations. A strong spatial signal will be required to improve locations with small error, but even a weak or complex signal in species distributions may improve very inaccurate locations. For our simulated data with strong spatial signals relative to size of the modelled domain, the mean distance error was around 15% of the domain size. For our real data and models fit using nine fish species, the location of fishing was identified to within 20-100 km, in a fishing region around 600 km in length. This fishery currently reports locations on a grid with a ∼10 km resolution, so these models would only be useful if the reported grid cells were considerably wrong. However, these models would improve on the prior reporting grid with 100 km resolution. The fishing domain in this trawl fishery is relatively small, and it’s likely that much larger domains would benefit relatively more from spatial signals of the scale seen here.

### 4.1. When to use which approach

When a relatively small subset of the location data is uncertain, perhaps less than 50% of the total, then the location error approach is effective and has neat statistical properties. This approach is essentially modelling location as a noisy variable, but treating only some locations as noisy and estimating (rather than conditioning on) that noise. However, for tasks that require the estimation of many uncertain locations, it makes more sense to fit a model using only accurate observations and then use that model for prediction.

If prior information is uncertain or weak, and the spatial signal in species distributions is weak, the location as response approach seems most effective. It was challenging to specify a useful prior in these cases, even when the true data were known (Fig. 4). The random forest is a particularly fast and effective model for this approach, and requires much less statistical input by the user. The modelling direction is illogical which could cause issues, but machine learning is an obvious choice when there are many observations to estimate (Watson et al 2023). In cases where location error is smaller, or priors are well informed, the inverse prediction approach appears useful and accurate. The inverse prediction approach has the advantages of a logical ‘causal’ modelling direction, and the addition of priors to help constrain the catch data. For example, if we know that a fishing event occurred in a grid cell, we can use that as a prior and find the maximum likelihood only in that cell (Fig. 6). This could pose a challenge when using the random forest, which cannot provide a location estimate if the predicted location (and error) is outside that specific cell.

**Fig. 6.**
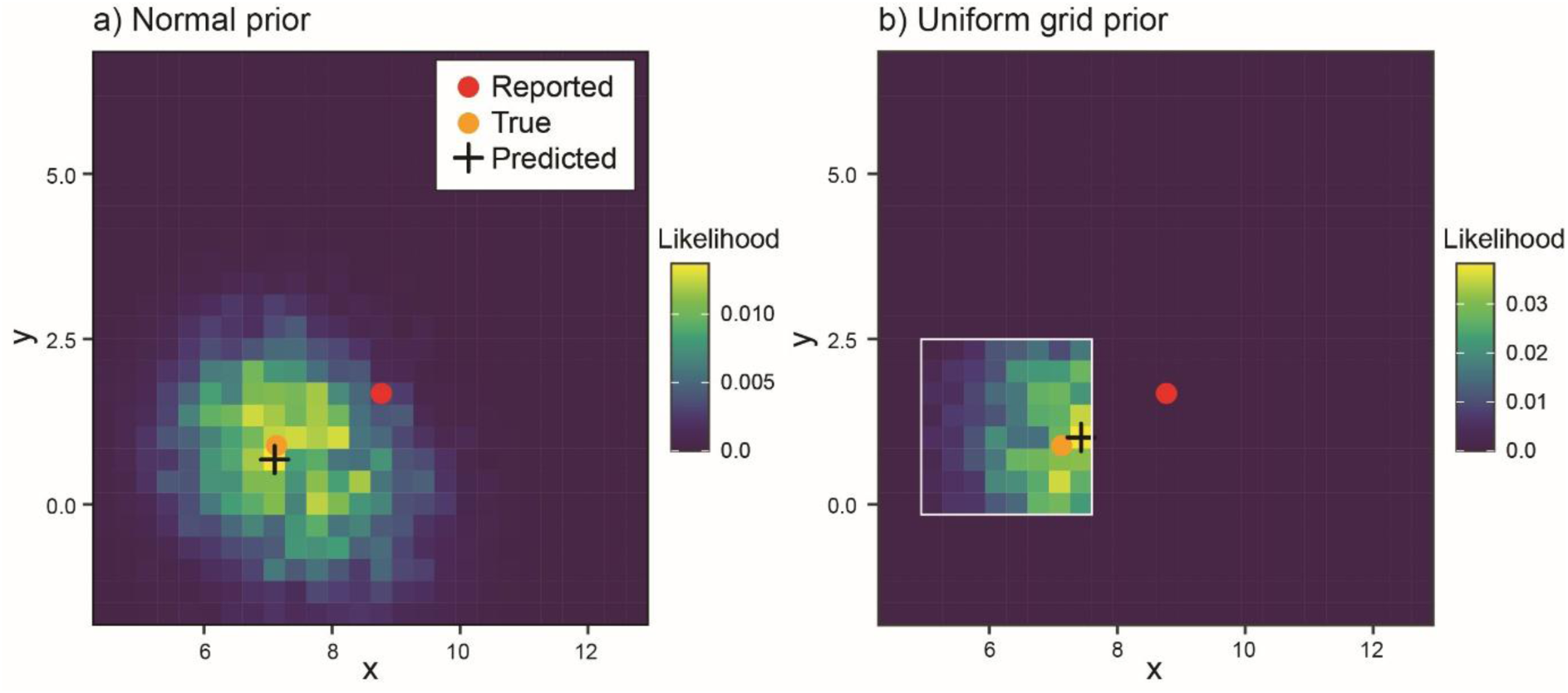
Examples of the posterior surface for the inverse prediction approach given a normal prior (a), or a uniform prior within a large grid cell (b). The latter can occur when reporting requirements are for a coarse grid (the white square in b) and we have no prior knowledge of where fishing occurred within that grid cell. This example has the same settings as Scenario 1 (Fig. 4).

### 4.2. Additional developments and caveats

There are many ways to construct SDMs, especially those modelling multiple species. Although modelling dependence among species can be valuable in joint modelling (Ovaskainen and Abrego 2020) the improvement to prediction can be small (Zurell et al 2020, Smith and Johnson 2025). Additional features such as estimating residual species correlations were ignored to avoid unnecessary complexity, although the location error and inverse prediction approaches can be extended to include these features. Also, in the real data scenario, a subset of reasonably common species with strong spatial signals we pre-selected to reduce the computational burden, which reduces the value of joint species modelling.

If the location at which a species assemblage is observed is uncertain, then the typical predictors used to build SDMs such as temperature are likely also uncertain (in reality, these variables will often have observation error but this is usually ignored). In the case of location estimation modelling, there are a few options to deal with uncertainty in spatially-structured predictors: 1) use best estimates for these additional predictors but acknowledge these sources of inaccuracy; 2) use a variety of likely values to explore sensitivity of location estimates to this variation; 3) leave these uncertain predictors out with the assumption that the species composition can recreate much of the signal; or 4) model these too as latent variables to estimate the observation error. The final option would be nicely integrated but computationally challenging.

An issue with inverse problems (such as the inverse prediction approach) is that small errors in observation can cause large uncertainty in the solution, although priors help stabilize the problem (Latz 2023). This helps explain why the inverse prediction approach is best used with informative priors to refine uncertain locations rather than estimate unknown ones. The choice of species could also lead to instability, which suggests some pre-filtering of species is beneficial. It would also be valuable to explore how the inverse prediction approach can be improved using machine learning algorithms (Kamyab et al 2022).

## Conclusion

These three approaches are all useful for estimating location from species composition, each with certain advantages. For simplicity and accuracy, the location as response does seem ideal for exploration and prediction. And as seen in other SDM research (Stock et al 2020, Smith and Johnsn 2025), random forests are a powerful method that should be included in the modelling toolbox. Our study shows that poor location accuracy can be substantially improved based on species compositions, especially given a set of accurate locations. This enhances goals such as the detection of illegal or misreported fishing (Watson et al. 2023), spatial management, and ecological monitoring, and could improve data quality in diverse applications such as citizen science surveys or historical records.

## Acknowledgements

This research did not receive any specific grant from funding agencies. We are grateful to the fishermen and researchers who contribute to the observer program.

## Data availability

R code to generate the simulated data and analyse it are shared on GitHub: https://github.com/smithja16/Location_From_Species. The scientific observer data are considered confidential data by the NSW Government and cannot be shared.

